# Epidemic network analysis for mitigation of invasive pathogens in seed systems: Potato in Ecuador

**DOI:** 10.1101/107367

**Authors:** C. E. Buddenhagen, J. F. Hernandez Nopsa, K. F. Andersen, J. Andrade-Piedra, G. A. Forbes, P. Kromann, S. Thomas-Sharma, P. Useche, K. A. Garrett

## Abstract

Seed systems have an important role in the distribution of high quality seed and improved varieties. The structure of seed networks also helps to determine the epidemiological risk for seedborne disease. We present a new method for evaluating the epidemiological role of nodes in seed networks, and apply it to a regional potato farmer consortium (CONPAPA) in Ecuador. We surveyed farmers to estimate the structure of networks of farmer seed tuber and ware potato transactions, and farmer information sources about pest and disease management. Then we simulated pathogen spread through seed transaction networks to identify priority nodes for disease detection. The likelihood of pathogen establishment was weighted based on the quality and/or quantity of information sources about disease management. CONPAPA staff and facilities, a market, and certain farms are priorities for disease management interventions, such as training, monitoring and variety dissemination. Advice from agrochemical store staff was common but assessed as significantly less reliable. Farmer access to information (reported number and quality of sources) was similar for both genders. Women had a smaller amount of the market share for seed-tubers and ware potato, however. Understanding seed system networks provides input for scenario analyses to evaluate potential system improvements.

---

Networks of crop seed distribution are an important factor in determining the success of agricultural systems. They drive the spatial distribution of crop plant genotypes and disease resistance genes, as well as the spread of seedborne disease. Seed systems encompass biophysical elements as well as all the stakeholders and activities that support the system, including interacting scientific (e.g., breeding, extension), management (e.g., agricultural practices, integrated pest management) and regulatory components (e.g., legally certified seed standards; Almekinders et al. 2007; Devaux et al. 2014; Jaffee et al. 1992; Kromann et al. 2017; Thiele 1999; Thiele et al. 2011). Thus, seed systems are best understood as a network of interacting biophysical and socioeconomic elements (Leeuwis and Aarts 2011). Establishing new seed systems has often been challenging, especially in low-income countries, probably in part due to the many system components that must dovetail for seed system success. We propose a framework for improving understanding of epidemiology in seed systems, taking into account socioeconomic components.

Ideally seed systems give farmers access to affordable disease free, disease resistant, high quality seed. In practice, most farmers in low-income countries (e.g., 98% of potato farmers in the Andes) save seed from the previous season for replanting (Devaux et al. 2014; Jaffee et al. 1992). Farm yields using saved seed are often poor compared to those obtained when using “improved seed”, obtained through integration of enhanced on-farm management, including disease resistance deployment, along with certified seed use as warranted. The recommended suite of practices for seed system enhancement has been proposed as an “integrated seed health strategy” (Thomas-Sharma et al. 2017). Scientists contribute to seed systems by developing more disease resistant varieties with other positive traits for dissemination through the system. Understanding seed systems can help scientists develop recommendations for system improvement based on linked epidemiological patterns and socioeconomic factors across a range of scales.

The risk of seedborne disease is particularly important in vegetatively propagated crops, such as potato, sweetpotato, yams, cassava, banana, and grafted fruits, compared to “botanical seed” or “true seed”. “Seed degeneration” is the reduction in yield or quality caused by an accumulation of pathogens and pests in planting material over successive cycles of vegetative propagation (Thomas-Sharma et al. 2017; Thomas-Sharma et al. 2016). Epidemiological models for vegetatively propagated crops must take into account the accumulation and spread of disease in planting materials (Thomas-Sharma et al. 2017). While seed transaction networks are sometimes studied and characterized (Labeyrie et al. 2016; Poudel et al. 2015; Ricciardi 2015; Tadesse et al. 2016; Violon et al. 2016), there is great potential for developing new approaches to predict the spread of seedborne diseases and help target disease detection efforts, training, treatments and other interventions (Andersen et al. 2017; Hernandez Nopsa et al. 2015; Pautasso et al. 2013; Tadesse et al. 2016). Here we use epidemiological network analysis (Shaw and Pautasso 2014) of a seed potato network to understand and predict disease risk, to develop a new type of scenario analysis for interpreting epidemic risk in seed systems that takes into account farmer information sources.

Efforts to improve seed systems often fail to improve the disease status of crops (Devaux et al. 2014; Devaux et al. 2010; Hirpa et al. 2010; Jaffee et al. 1992; Kromann et al. 2017; Panchi et al. 2012; Thiele et al. 2011; Thomas-Sharma et al. 2016). Understanding the structure and function of formal (state regulated systems (Sperling et al. 2013)), informal, and mixed seed systems can support the development of more sustainable seed systems. Aspects that determine the degree of seed system utility, sustainability, and resilience include access to and availability of seed, seed quality, cultivar quality (e.g., adapted, disease resistant, and matching user preferences), affordability, and profitability (Sperling et al. 2013). There are tradeoffs in connectivity for farmers, where high connectivity is good for getting access to new varieties and training, but can increase the risk of being exposed to disease. Managing connectivity can help to increase system resilience (Biggs et al. 2012).

Seed system resilience is tested when there are significant stressors or crises, be they environmental (Violon et al. 2016), biotic (e.g. pathogen or pest outbreaks) or socioeconomic (McGuire and Sperling 2013). Though broad categories of threats are predictable, some events may be viewed as crises because they are spatially varied, temporally unpredictable, and may have multiple distinct drivers (e.g. pathogen, drought, conflict and economic crises). In high-income countries, regulation plays a substantial part in keeping a profit-driven sector functioning in everyone’s interests (Frost et al. 2013). However, formal seed systems can be “static and bureaucratic” (Lybbert and Sumner 2012) where seed certification standards are unachievable with reasonably available resources. Often resource-poor farmers are priced out of the formal system, or government subsidized systems can be unreliable. Sometimes improved varieties require inputs that are out of reach of resource-poor farmers, or disease pressure is enough to require many inputs. These are common explanations for the persistence of lower performing informal seed systems even after interventions that seek to improve them. Given the long persistence of informal seed systems, it appears that single optimal solutions are unlikely. Often governmental and aid based interventions emphasize provision of certified clean seed of traditional and improved varieties to as many resource-poor farmers as possible (Tadesse et al. 2016). Typically, such seed systems revert to informal ones where farmers use their own seed. Such interventions are repeatedly attempted, suggesting that changes to on-farm disease management practices might provide comparable yield benefits (Thomas-Sharma et al. 2017) while being more sustainable within persistently informal systems. Despite repeated failures, development agencies continue to orient their interventions toward the development of regulated (McGuire and Sperling 2013; McGuire and Sperling 2008) demand-driven systems that support a for-profit model of seed supply, believing them more sustainable and resilient (McGuire and Sperling 2013; Sperling et al. 2013). Often, after project funds are discontinued, the subsidized formal seed systems revert to largely informal ones with poor access to improved seed. A common belief is that this lack of resilience relates to a lack of diversity in terms of crops and cultivars, or supply channels (McGuire and Sperling 2013). Given a long history of aid to improve seed systems it would seem that farmer decision-making is poorly understood.

A frontier for plant epidemiology is to better incorporate and model disease spread while taking into account actual human decision making about disease management (McRoberts et al. 2011). Seed system development efforts often attempt to foster equitable access by stakeholders to services (Ricciardi 2015), although more needs to be understood about the effects of gender and other individual traits on access. Epidemiological network analyses can help to identify systemic vulnerabilities related to gender access to quality planting materials, integrated pest management information, and the market for products (Tadesse et al. 2016). Clearly short- and long-term planning by government agricultural agencies, farmers, and aid agencies could help to meet the variety of seed supply challenges. Stakeholders, especially governments and non-governmental agencies need to be flexible to strike a good balance between sustaining and transforming systems. Trade-offs are likely, with interventions under one scenario or set of stressors potentially being counter indicated in another scenario, or for some stakeholders.

The risk that pathogens can move through a seed system network is a key component of disease risk, along with other risk factors such as potential transmission by vectors or wind dispersal. Detection of pathogens in a seed network in a timely manner can allow for mitigation measures to be implemented. Hub nodes (nodes with many links) and bridge nodes (nodes that connect distinct regions of a network) will tend to have important roles in the risk of disease spread, and in sampling and mitigation (Hernandez Nopsa et al. 2015). However, nodes on the periphery of a network could be the entry point for an invasion of that network (Xing et al. 2017). While the importance of hub and bridge nodes is intuitive, key roles of other nodes may be revealed in more detailed analyses of likely patterns of disease spread. Strategies for dissemination of resistant varieties may need to change depending on network properties. In addition, the spread of endemic pathogens such as *Rhizoctonia* spp., or the potential arrival of emerging diseases from distant locations (e.g., *Dickeya* spp.; Czajkowski et al. 2015; Czajkowski et al. 2011; van der Wolf et al. 2014) can be modeled and mitigation strategies tested using a multilayer network analysis (Garrett 2012, 2017).

Exponential random graph models (ERGMs) can be used to characterize networks in terms of the likelihood that links exist between different types of nodes (Handcock et al. 2008). ERGMs have been used extensively in social sciences, and can be used to identify actors that have key roles in epidemics or experience particular risk. In plant disease epidemiology, ERGMs have the potential to contribute to analyses of human effects on and responses to disease risk, and of interactions among different types of pathogens, vectors, and environments (Welch et al. 2011).

The study presented here addresses the challenge of understanding the strengths and vulnerabilities of multilayer seed system networks, considering both the network of seed transactions and the network of communication about IPM. We introduce a new type of scenario analysis for studying potential epidemics in seed transaction networks, and the role that particular network nodes play in sampling and mitigation of epidemics. This analysis focuses on the component of disease risk due to the structure of seed networks. To this end our objectives are to (a) characterize cultivar dispersal through a potato seed system in Ecuador; (b) determine whether gender is associated with different types of network transactions or access to information; (c) model the potential spread of a seed borne pathogen through the seed system in order assess the risk level at each node, to evaluate their utility as control points for pathogen mitigation measures; and (d) characterize how the seed system transaction network might adapt to a scenario where the CONPAPA management team and consortium no longer plays an organizing role, and existing seed multipliers must compensate for its absence.

## MATERIALS AND METHODS

### Study system context: seed degeneration

Viruses such as *Potato virus Y* (PVY), *Potato virus X* (PVX) and *Potato leafroll virus* (PLRV), are major causes of seed degeneration in many parts of the world (Frost et al. 2013; Salazar 1996). Additionally, depending on the geographic region, fungi, bacteria, nematodes, phytoplasmas, and insects can also play important roles in potato seed degeneration (Thomas-Sharma et al. 2016). In high-elevation potato production regions of Ecuador, *Rhizoctonia solani* is a major cause of seed degeneration (Fankhauser 2000), while in many other tropical and subtropical countries *Ralstonia solanacearum* is a major concern (Mwangi et al. 2008). Adding to this complex etiology, the rate of degeneration is also highly variable across geographical regions. Factors such as host physiology, vector dynamics, environmental variability, and the choice and success of management strategies can affect the rate of degeneration (Thomas-Sharma et al. 2017; Thomas-Sharma et al. 2016). In high elevation regions, for example, lower temperatures can limit vector activity and pathogen multiplication while also influencing host physiology that limits pathogen transmission into daughter tubers (Bertschinger 1992; Navarrete et al. 2017). In at least one case the presence of *Potato yellow vein virus* (PYVV) was associated with small yield *improvements*, possibly via some sort of competitive interaction with other viruses (Navarrete et al. 2017). In the Andes, evidence suggests virus transmission to daughter tubers is usually incomplete with between 30 and 75% of tubers being infected (Bertschinger et al. 2017). Similarly, the application of management strategies such as resistant cultivars, certified seed material and other on-farm management strategies, individually and/or collectively, can affect the spread of disease epidemics in a region (Thomas-Sharma et al. 2017). A better understanding of these inter-related factors could contribute to the design of an integrated seed health strategy for a geographic region (Thomas-Sharma et al. 2016).

### Study system: the CONPAPA potato seed system in Tungurahua, Ecuador

There are approximately 50,000 ha of potato production in Ecuador, with 97% of this area located in the Andes, and 87% of farms being less than 10 ha in size (Devaux et al. 2010). It is possible to produce tubers all year, which has created a market that expects fresh potatoes for consumption year round (Devaux et al. 2010). Seed tubers from a farmer’s previous season are typically planted in the next. This makes the potato crop subject to seed degeneration and yield losses (Thomas-Sharma et al. 2016). The national agricultural research institute, INIAP (*Instituto Nacional de Investigaciones Agropecuarias*) is the only agency in Ecuador registered to produce formal basic seed potato. However, according to a 2012 estimate, less than 3% of the seed potato used in Ecuador is from the formal system (Thomas-Sharma et al. 2016). Two preferred cultivars for farmers in the Ecuadorian Andes are INIAP-*Fripapa* and *Superchola*. However, farmers also grow many other cultivars, such as INIAP-*Gabriela*, INIAP-*Catalina*, and *Diacol-Capiro*. Seed is produced by INIAP from pre-basic seed, which are mini-tubers produced from *in-vitro* plants. Basic seed, the next generation, is multiplied in the field by INIAP or associated farmers. The next three generations of seed include the following three seed categories; registered seed (*semilla calidad I*), certified seed (*semilla calidad II*), and selected seed (*semilla calidad III*), and are produced in the field by seed producers. Trained seed producers form a part of the Consortium of Potato Producers (CONPAPA) and produce seed for member farmers (Fig. 1). The yield increase associated with each of these three categories has been reported to be 17%, 11% and 6%, respectively, compared to the seed produced by the farmers in the informal system (Devaux et al. 2010), although these estimates are low compared to the potential (30%) yield increases reported globally from the use of quality seed potato (Thomas-Sharma et al. 2016). Established in 2006, CONPAPA has a membership of ca. 300 farmers in central Ecuador (principally in Tungurahua, Chimborazo and Bolívar Provinces). This organization is the current realization of various aid and governmental efforts to improve livelihoods for small-scale potato farmers (Kromann et al. 2017). It aims to support small-scale farmer associations that produce seed potato and potato for consumption (ware potato), through training, provision of quality assessed seed, and by processing and marketing produce. It cleans and processes produce (e.g., for chips and fresh potato) in regional processing facilities. It also sells potato on behalf of members. Annual mean, production yield of ware potato in CONPAPA (Tungurahua) ranges between 15 and 20 metric tons per hectare, with production levels being influenced by management, variety, time of year, and the number of generations since the seed was sourced from basic seed. Average production reported by CONPAPA is higher than the 9.5 metric tons per hectare that has been reported for Ecuador as a whole (Devaux et al. 2010). CONPAPA in Tungurahua reported (www.conpapa.org) that it supplies more than 25 tons of potato for consumption per week to meet market demand (0.3% of Ecuador’s total production; Devaux et al. 2010). Importantly, CONPAPA has trained seed multipliers who provide seed for redistribution to member farmers.

**Fig. 1.**
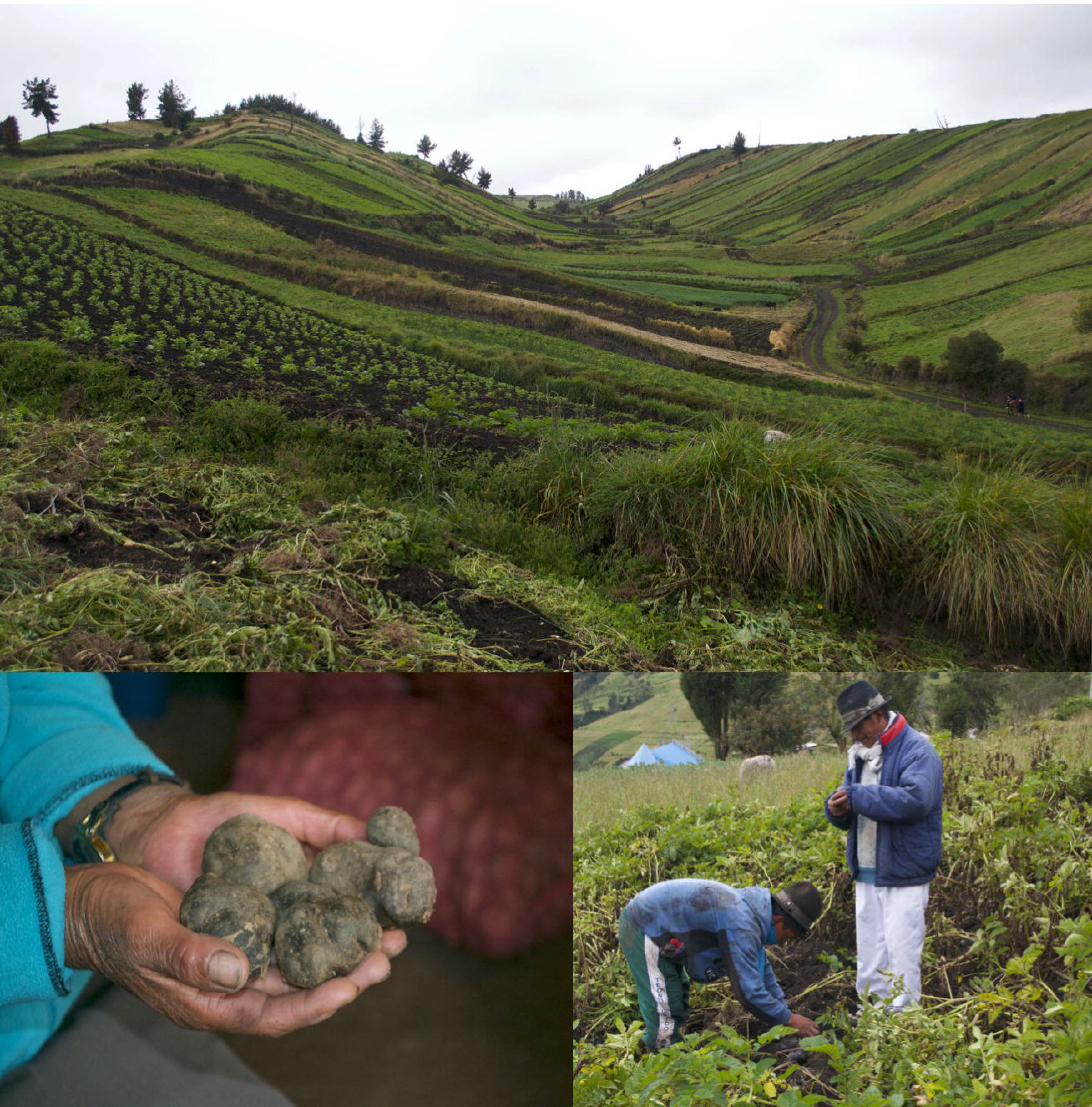
Potato production by farmers in the CONPAPA seed system in Tungurahua Province, Ecuador (photos: J. F. Hernandez Nopsa).

### Survey methods

This study focuses on 48 farmers who are members of CONPAPA in the Tungurahua province. This is 66% of the 72 heads of households registered as members in this region (Montesdeoca, pers. comm.). However, the 48 farmers in this study represent a census of all the active farmers at the time of this study. (We conceptualize the farmers’ reported transactions as a sample of their typical types of transactions across seasons.) Farmer network sizes and farmer activity can change as farmers opportunistically pursue a variety of alternative livelihoods from year to year, e.g., construction or service jobs, in response to changing conditions (in good and bad years; Violon et al. 2016). A survey was completed by scientists via on-farm voluntary interviews of 48 farmers in the CONPAPA district of Tungurahua over three weeks in November and December, 2015. In addition to demographic information, the survey documented the seed sources, cultivars planted, volume bought, and price paid for the last three planting periods, as reported by farmers. Farmers were also asked to report (a) the sale or use of potato for food, including destination, cultivar, volume, and price received, (b) the principal pests and diseases they observe, and (c) their sources of advice regarding integrated pest management, and the confidence they had in that advice. In some cases, there was missing data related to volume or price information. The results of this survey are available at http://dx.doi.org/10.21223/P3/XKHUTL and the data specifically used in these analyses are included as supplemental material.

### Data analysis and modeling

Networks of seed and ware potato transactions between the farmers and other stakeholders were analyzed using the igraph and statnet packages (Csárdi and Nepusz 2006; Handcock et al. 2008) in the R programming environment (R Core Team 2017). Selected R scripts and the resulting output are available through links at http://www.garrettlab.com/epid-seed/, along with an interactive interface for understanding the structure of epidemic risk in the CONPAPA system. The adjacency matrix we evaluated was based on reported sales, where a link indicates a directed transaction resulting in the movement of seed or ware potatoes. For cases where farmers reported a transaction but did not give volume information, links were depicted in the network as dotted lines and given a minimum visible width. Missing price and volume data were not treated as zeroes, but were omitted from the calculation of means and percentages. Missing volume and price data are reported in the results. Transaction counts, volumes and prices were compared with respect to potato cultivar and farmer gender, based on percentages, means, and two-sided Wilcoxon tests (using the wilcox.text function in R). While the farmers sampled represent a complete census of the CONPAPA farmers, we treat their reported information as a sample of reported transactions across years. We also evaluated the effect of node type (farmer or institution) on the likelihood that a link exists in the potato transaction network using an ERGM in the statnet package in R (Handcock et al. 2008).

We also evaluated a second adjacency matrix describing communication, based on the information sources that farmers reported related to disease and pest management. Links in this matrix indicate the reported flow of information. Based on the structure of this network, the information access of each node (farmer and/or other information source) was evaluated in two ways. “Information quantity” was defined as the number of information sources a node accesses, or the in-degree for a node. “Information quality” was defined as the maximum level of trust reported for any of a node’s sources of information.

The frequency with which common pests and diseases were reported by farmers, including diseases responsible for seed degeneration, is reported overall and by gender (where gender differences were tested using chi-square tests).

### Rating the importance of nodes for sampling efforts

Optimal management of potential invasive pathogens in a seed system depends on identifying the most important geographical nodes for sampling to detect disease (both in the field and in the harvested tubers). Sampling some nodes will tend to result in rapid detection of the pathogen, while sampling other nodes will likely only detect the pathogen after it has already spread widely in the network. In a scenario analysis, disease spread was simulated across the seed and ware potato distribution network, where the network was based on reports aggregated across the last three plantings (and actual or anticipated harvest dates ranged from May 2014 to May 2016). In the simplest version of the analysis, each node was considered equally likely to be the point of initial introduction of a pathogen into the seed system network. Another version of the analysis drew on the structure of the communication network. In this case, the probability that a pathogen would be introduced into the network by a given farmer was weighted by a function of that farmer’s level of information quantity or quality (defined above), as a proxy for the node’s ability to respond effectively. The idea is that a well-informed farmer (with high information quantity and/or quality) will be less likely to be a point of disease introduction into the network, and will be more prepared to keep a new pathogen from becoming established.

Information quantity and quality were transformed to values that act as proxies for the likelihood that a node does <u>not</u> effectively manage an invasive pathogen, because of inadequate information. For information quantity, we considered the probability (p_1_) that the necessary information is <u>not</u> obtained from a given source node. Information quantity was transformed as the probability that the information was not received from <u>any</u> of the potential sources, as p_1_ to the power of the node in-degree, where the results were evaluated for p_1_ = 0.1, 0.5, and 0.9. (For nodes for which we had no reports about degree, where the results were evaluated for p_1_ = 0.1, 0.5, and 0.9. (For nodes for which we had no reports information quantity, in-degree was set to 3 for individuals, 10 for institutions, and 0 for markets.) Information quality was sampled as a reported level of trust (y) for each information source, on a scale of 0 to 5. Information quality was transformed as 1 – max(y)/5. Because max(y) was usually 5, we scaled this back to consider scenarios without “certain successful management”, by multiplying 1 – max(y)/5 by 0.1, 0.5, and 0.9. (For nodes for which we had no reports about information quality, max(y)/5 was taken as the reported farmer average, 0.9, for nodes representing other farmers, as 1 for nodes representing institutions, and as 0 for markets.)

The simulation of epidemic spread generates an estimate of the number of nodes infected before the disease will be detected at each potential sampling node, given that each potential starting node has a weighted probability of being the initial source based on information quantity or quality. The output allows us to estimate relative risk in terms of the number of nodes that would be infected if only the node in question were monitored.

### Scenario analysis where the CONPAPA management team does not supply seed

The CONPAPA management team is clearly central to this seed system, a key “cutpoint”, or node whose removal creates multiple disconnected components in the network. We explore how resilient the seed system might be if the CONPAPA management team were removed. How would other nodes need to compensate for its absence? We compared the scenario where the CONPAPA management team provides seed to farmers and multipliers with a scenario where it does not have a role in seed provision. For this alternative scenario we evaluated the reported volumes for seed transactions over three plantings. Then where the CONPAPA management team provided basic seed to multipliers we replaced these transactions with INIAP, the government agency that provides basic seed to CONPAPA (GovtAgency1 in the Figures). Finally, where farmers sourced their seed from the CONPAPA management team, they instead sourced their seed from the geographically nearest multiplier (Farmers 7, 27, 34 and 46). The alternative scenario thus maintains the same transaction volumes that were reported but removes the CONPAPA management team as the go-between replacing these with the most plausible alternative. We evaluate the structure of this new network.

## RESULTS

### Seed system: overview

The seed system network depicted in this study is sparse, has highly heterogeneous in-degree, with a degree of clustering and higher-level cycles, while links are directed, weighted, and dynamic. It is centered around the CONPAPA management team in Tungurahua, which provides and receives seed and ware potato from member farmers (Fig. 2). A total of 1157 quintals (45.36 kg bags), or 52 t (metric tons), of seed was reported as used by farmers in the most recent planting, where CONPAPA provided 47%, 36% was farmers’ saved seed, and the remaining 16% came from other sources. CONPAPA was reported as receiving only 7 t of seed from trained male seed multipliers. Only two women (F7 and F46) reported providing seed (Puca, Fripapa and Superchola) to CONPAPA during this interval, although farmers 7, 8, 10, 19, 36, 40, 46, and 47 are women trained to be seed multipliers. Of the 48 farmers that reported buying or selling potato or seed, 16 (33%) were women. Farmers reported a total of 503.9 *t* potato being sold, with CONPAPA buying 414.7 t (82%) of potato (where 28% of this was from women). Farmers reported selling 85.3 *t* directly to local markets, and one farmer reported selling 3.2 t directly to a restaurant. It is important to note that 262 transactions were reported in the most recent season but interviewees did not provide volume for 71 transactions or price information for 58 transactions (including self-supplied seed transactions). On a per transaction basis there was a difference between the volume of ware potato sold by women (median=5 quintals) and men (median=40 quintals; Wilcoxon test (two-sided), W = 2594, p-value = 1.8 e-08). In the ERGM analysis, node type had an effect on the likelihood that reported links exist in the seed and ware potato transaction network (p < 0.0001). Farmers were much more likely to report transaction links with institutions than with other farmers, indicating that the system is “formalized” to a great extent. There was also evidence for a difference in per transaction volume for seed tubers between women and men, with medians of 3 and 5 quintals respectively (Wilcoxon test, two-sided, W = 5142.5, p-value = 0.0002). There was no evidence for a difference in per transaction prices for ware potato for women and men, with medians of 15 USD per quintal being received by both genders (Wilcoxon test (two-sided), W = 2513, p-value = 0.9). Prices were infrequently reported. Unreported here is the movement of pre-basic seed to CONPAPA from INIAP. CONPAPA in Tungurahua may also receive seed from CONPAPA multipliers outside of the region. Farmers reported replacing seed every 3-4 seasons, indicating that purchased seed is grown alongside seed saved from previous plantings.

**Fig. 2.**
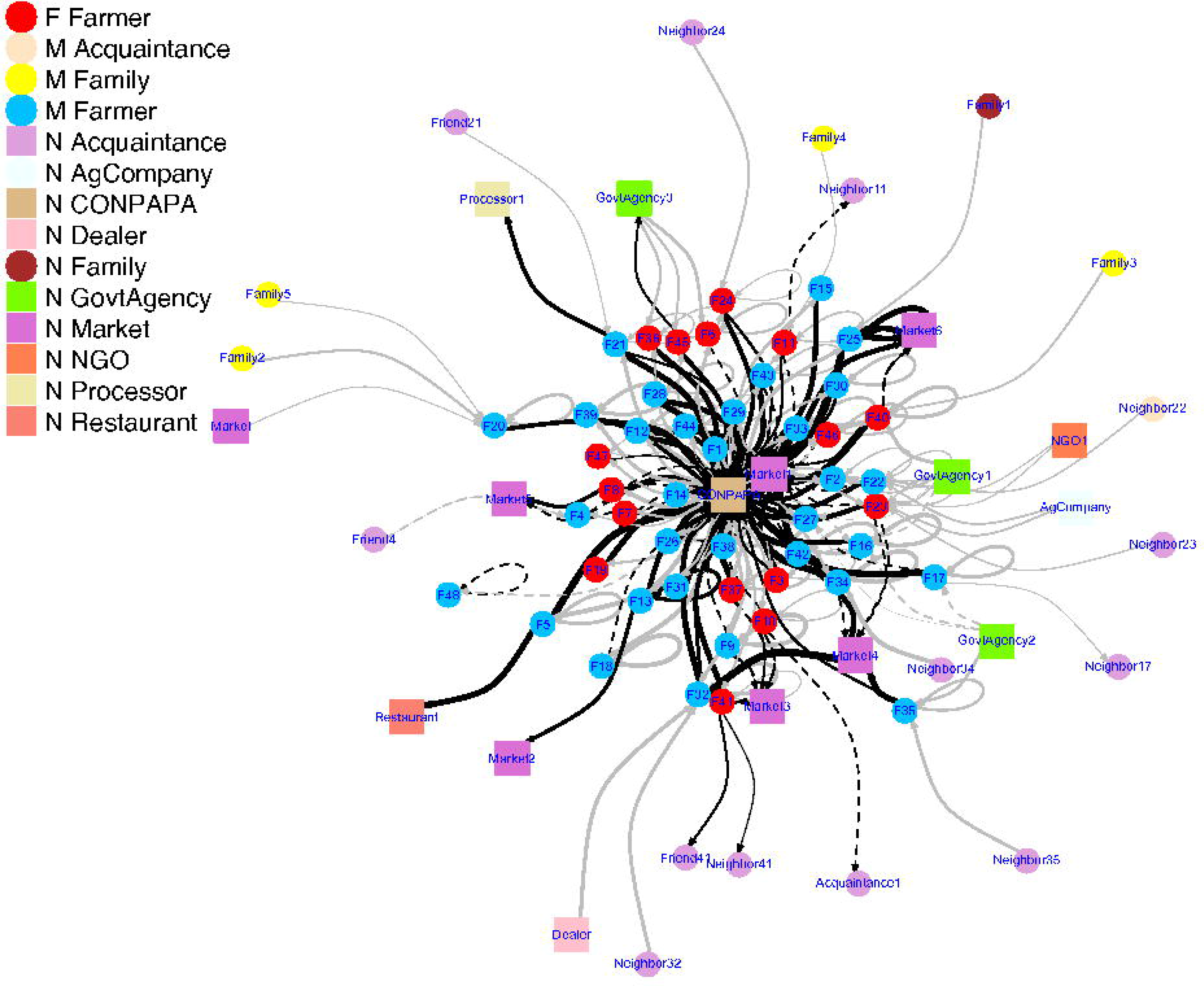
A seed system transaction network in which nodes represent 48 farmers associated with CONPAPA in Tungurahua, Ecuador, along with other institutions and individuals linked with them. Links indicate potato movement, and are weighted by the volume (proportional to line thickness) of seed potato and ware potato bought, sold, used or traded by farmers. Data are from the three most recent seasons reported in November 2015. Black lines indicate seed, and gray lines represent potato for food consumption. Self-loops represent seed produced on-farm. Dotted lines represent transactions where volumes were not reported. (The Fruchterman Reingold layout was used for generating the network representation in this and subsequent figures.)

Most transactions were with CONPAPA and the market in the nearby town of Ambato. More than 88% of the transactions were between actors that were >10 km apart. There were no CONPAPA member farmer to farmer transactions. Only 7% (40) transactions were with neighbors and 1% were with family members (for which there was no geographic location data).

### Seed system: analysis by variety

Overall, while farmers planted on average two cultivars, the median use was just one. In other words, about half of the farmers reported planting a single cultivar, while the other half planted 2 to 5 different cultivars. Ranking the use of cultivars by the numbers of farmers using them matches almost exactly the ranking by number of transactions per cultivar (Table 1), which suggests that the high number of transactions for the main cultivars is driven by their overall popularity. The three most commonly planted cultivars, according to these criteria, were Superchola (33% of farmers planted it, its product transactions represent 36% of all transactions, and its seed transactions 32%), Fripapa (17%, 20%, 22%) and Puca (13%, 10%, 10%), in respective order of ranking.

**Table 1.** The number of transactions and number of farmers using each cultivar for the current season, as reported in November 2015 (with percentages).

| Cultivar | Total transactions | Seed potato |  | Ware potato |  | No. Farmers using the cultivar |  |
| --- | --- | --- | --- | --- | --- | --- | --- |
| Superchola | 90 | 40 | 32% | 50 | 36% | 31 | 33% |
| Fripapa | 56 | 28 | 22% | 28 | 20% | 16 | 17% |
| Puca-shungo | 27 | 13 | 10% | 14 | 10% | 12 | 13% |
| Yana-shungo | 17 | 9 | 7% | 8 | 6% | 8 | 8% |
| Unica | 16 | 7 | 6% | 9 | 7% | 6 | 6% |
| Carolina | 13 | 6 | 5% | 7 | 5% | 4 | 4% |
| Victoria | 10 | 4 | 3% | 6 | 4% | 4 | 4% |
| Gabriela | 8 | 3 | 2% | 5 | 4% | 3 | 3% |
| Chaucha | 7 | 4 | 3% | 3 | 2% | 3 | 3% |
| Carrizo | 6 | 3 | 2% | 3 | 2% | 2 | 2% |
| Suprema | 4 | 2 | 2% | 2 | 1% | 2 | 2% |
| Americana | 2 | 1 | 1% | 1 | 1% | 1 | 1% |
| Natividad | 2 | 2 | 2% | 0 | 0% | 1 | 1% |
| Norteña | 2 | 2 | 2% | 0 | 0% | 1 | 1% |
| Tulca | 2 | 1 | 1% | 1 | 1% | 1 | 1% |

A second comparison of the total volume of transactions by cultivar, shows that the three most frequently exchanged cultivars are also the ones with most transacted volume (Table 2). Indeed, Superchola’s transacted volume represents 40% of all volume transacted in terms of product and 35% in terms of seed. Fripapa’s seed volume transacted was higher than the product volume transacted at 26% vs. 21%. Finally, the volume of variety Puca represents 9% of ware potato and 7% of seed. Interestingly, two varieties that were not reported by the majority of farmers—Carrizo and Victoria—represented 8 and 7% in terms of volume transacted, almost as much as Puca. This occurred because a few farmers provided large volumes of product to a few non-CONPAPA buyers. Finally, the percentage seed volume transacted for Unica was larger than for Puca (9%) and Natividad was the same as Puca (7%).

**Table 2.** Volume of seed in quintals (where 1 quintal = 45.63 kg) and product exchanged (with percentages).

| Variety | Total volume |  |  |  | Volume per transaction |  |
| --- | --- | --- | --- | --- | --- | --- |
|  | product |  | Seed |  | Product | Seed |
| Superchola | 4580 | 40% | 425 | 35% | 143 | 11 |
| Fripapa | 2405 | 21% | 323 | 26% | 172 | 15 |
| Puca-shungo | 999 | 9% | 80 | 7% | 111 | 7 |
| Carrizo | 960 | 8% | 66 | 5% | 480 | 22 |
| Victoria | 760 | 7% | 48 | 4% | 190 | 12 |
| Unica | 600 | 5% | 116 | 9% | 200 | 19 |
| Carolina | 470 | 4% | 79 | 6% | 118 | 13 |
| Yana-shungo | 350 | 3% | 43 | 3% | 58 | 5 |
| Gabriela | 90 | 1% | 6 | 0% | 45 | 3 |
| Chaucha | 80 | 1% | 16 | 1% | 27 | 5 |
| Suprema | 15 | 0% | 21 | 2% | 15 | 11 |
| Americana | 0 | 0% | 1 | 0% | . | 1 |
| Tulca | 0 | 0% | 0 | 0% | . | . |
| Natividad |  | 0% | 90 | 7% |  | 45 |
| Norteña |  | 0% | 0 | 0% |  | . |

### IPM information

Farmers largely reported obtaining information about integrated pest and disease management (IPM) from the CONPAPA management team (mean in-degree for information received by farmers was 3.5 overall; Fig. 3). There was not evidence for a difference (Wilcoxon test, W = 236, p-value = 0.4668) between male (3.7) and female (3.2) in-degree with respect to number of information sources reported. Importantly, farmers frequently reported receiving information from agrichemical stores (green squares in Fig. 3). Family members also provided important sources of information about IPM (Fig. 3). A quarter of the women reported their husband as a source of information for IPM, but no men reported that their wife was a source of IPM information. Farmer assessed trust levels could range between zero and five. There was some evidence for a difference in median trust levels reported by men and women 3 compared to 5 for CONPAPA (Wilcoxon test, W = 5364, p-value = 0.02). Though the pattern was less obvious when trust levels women reported with respect to IPM information from their husbands was removed (median of 3.5, and 4 for men and women respectively, Wilcoxon test, W = 4825.5, p-value = 0.06). The main sources of information were CONPAPA and agrochemical stores, where the median trust level farmers reported for all stores was 3 compared to 5 for CONPAPA (Wilcoxon test, W = 457, p-value = 2.2e-0). Only one farmer reported the internet as an important source of information about management.

**Fig. 3.**
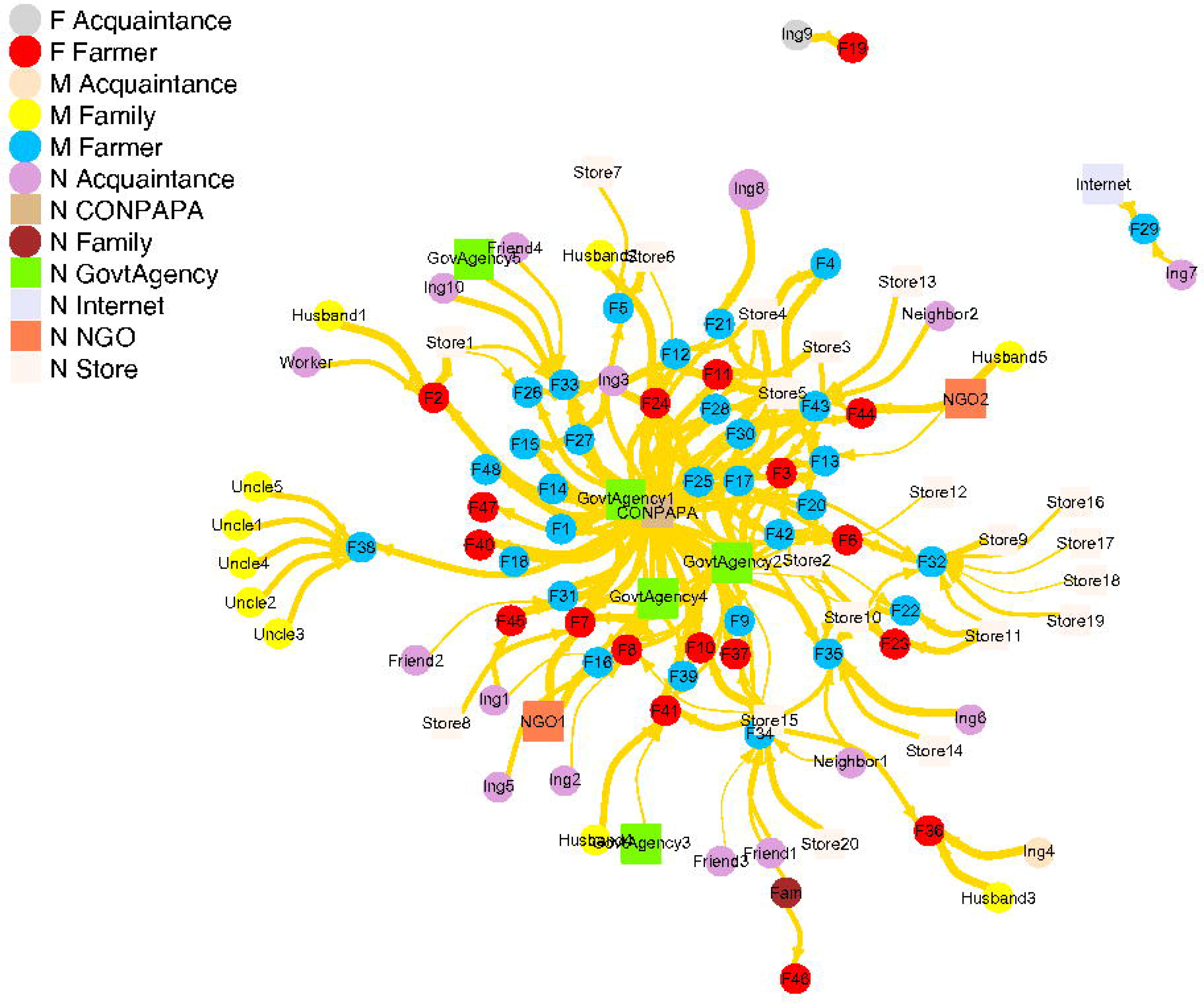
A network depicting farmer-reported information sources for integrated pest and disease management (IPM). Link thickness is proportional to the reported level of trust that the farmer has in that source of information.

The most frequently reported diseases and pests were potato late blight, Andean potato weevil, and potato black leg. Despite prompting, viruses were reported by only one percent of farmers (Table 3). Slugs and leaf miners were more frequently reported as a problem by women than men, though rates were low (Table 3).

**Table 3.** Pests and diseases reported by farmers in Tungurahua, Ecuador, in order by the frequency of reports. Pests and diseases known to cause seed degeneration are indicated.

| Pathogen/disease or pest | Causing degeneration | Women reporting | Men reporting | % farmers |
| --- | --- | --- | --- | --- |
| *Phytophthora infestans* (Late blight) | Yes | 15 | 30 | 94 |
| *Premnotrypes* spp. (Andean potato weevil) | Yes | 10 | 26 | 75 |
| *Rhizoctonia solani* (Potato black leg) | Yes | 7 | 16 | 48 |
| *Puccinia pittieriana* (Common rust) | No | 6 | 12 | 38 |
| *Epitrix* spp. (Potato flea beetles) | Yes | 3 | 9 | 25 |
| *Phthorimaea operculella*, *Symmetrichema tangolias*, and *Tecia solanivora* (Potato moths) | Yes | 4 | 4 | 17 |
| *Fusarium* spp. (Fusarium rot) | Yes | 1 | 6 | 15 |
| *Liriomyza* spp.* (Leaf miner) | Yes | 5 | 2 | 15 |
| Slugs* | No | 4 | 0 | 8 |
| *Frankliniella tuberosi* (Thrips) | Yes | 2 | 1 | 6 |
| Nematodes | Yes | 1 | 1 | 4 |
| *Spongospora subterranean* (Powdery scab) | Yes | 1 | 1 | 4 |
| *Septoria lycopersici* (Annular leaf spot) | No | 0 | 1 | 2 |
| Viruses | Yes | 1 | 0 | 2 |
| White fly | Yes | 1 | 0 | 2 |

Gender differences (\*) are significant in a Chi square test ( $\alpha=0.05$ , $df=1$ )

### Disease risk in the system

Under the scenarios we evaluated (Fig. 4A-C), the CONPAPA management team is the most effective place to monitor in order to detect a disease before it has spread far. This reflects its central role in the network. Similarly, several stakeholders and farmers at the periphery of the seed and potato network tend to be poor locations for detecting potential disease in every simulation. This is because they only provided seed rather than receiving seed or product (dark purple) in this network, or had low in-degree (blue or light purple; Fig. 4A-C). Weighting risk of establishment based on the information quantity for IPM (Fig 4 B) causes some nodes of intermediate importance to become more important for monitoring, particularly where we assume the first few information sources a node has access to have the greatest impact on management (p_1_ = 0.1). The results for the scenario with risk weighted as a function of the quality (trust) of information were very similar to the case where all nodes weighted equally. In all cases the market in Ambato, the largest town in the region, is also a good place to monitor, though we assume that even with good information available about IPM this would do little to change disease risk there. This particular market had the highest reported in-degree of any of the five markets. An important caveat is that clearly not all diseases can be mitigated effectively by IPM, this assumes that an IPM intervention is available to farmers so that they can reduce the probability of disease establishment.

**Fig. 4.**
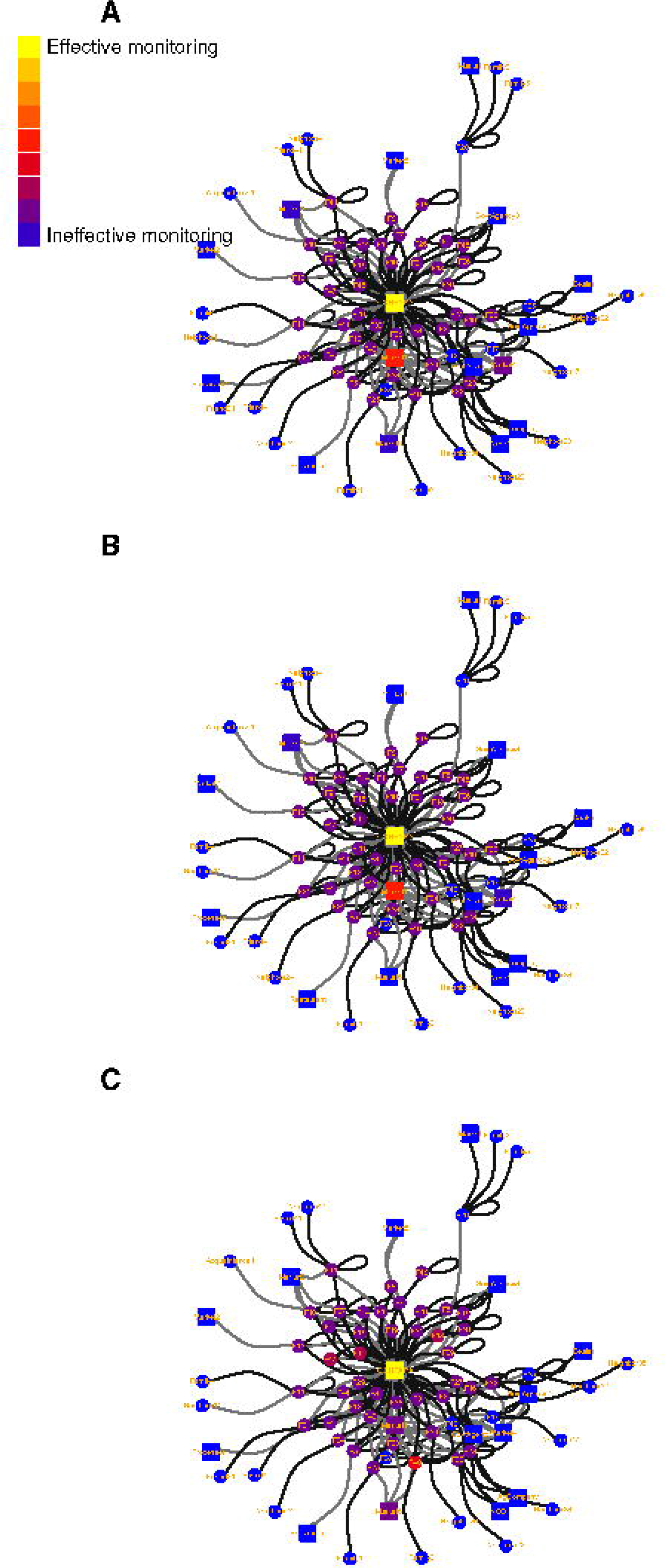
Pathogen invasion is simulated with initial infection starting at a random node and proceeding through the network defined by farmer transactions for seed (black) lines and ware potato (grey). Link widths are scaled to volume of transaction. This network represents the last three seasons reported in November 2015. The value of monitoring each node is evaluated in terms of the number of nodes (few=yellow, blue=many) that would become infected before the disease was detected at that node. Three scenarios were evaluated, where the probability/risk of the disease starting at a given farmer node is weighted differently. **A**, all farmers are equally likely to be an initial source of introduction of the pathogen into the network; **B**, risk of being an initial source is proportional to 0.9 to the power of the number of sources (node in-degree, not including self-loops) as depicted in the IPM information network in Fig. 3.; **C**, risk of being an initial source is proportional to 0.1 to the power of the number of sources.

### Scenario analysis where the CONPAPA management team does not supply seed

We compared the scenario where the CONPAPA management team provides seed to farmers and multipliers, and multipliers sell their seed to CONPAPA (Fig. 5A), with a scenario where the CONPAPA management team does not have a role in seed provision (Fig. 5B). In this analysis, based on a role of geographic proximity to multipliers, we see that multipliers do not have equal access to all the seed buying farmers in the market (Fig. 5B). CONPAPA’s role as distributor and organizer of seed distribution (Fig. 5A) may result in all farmers having access to seed from any of the multipliers.

**Fig. 5.**
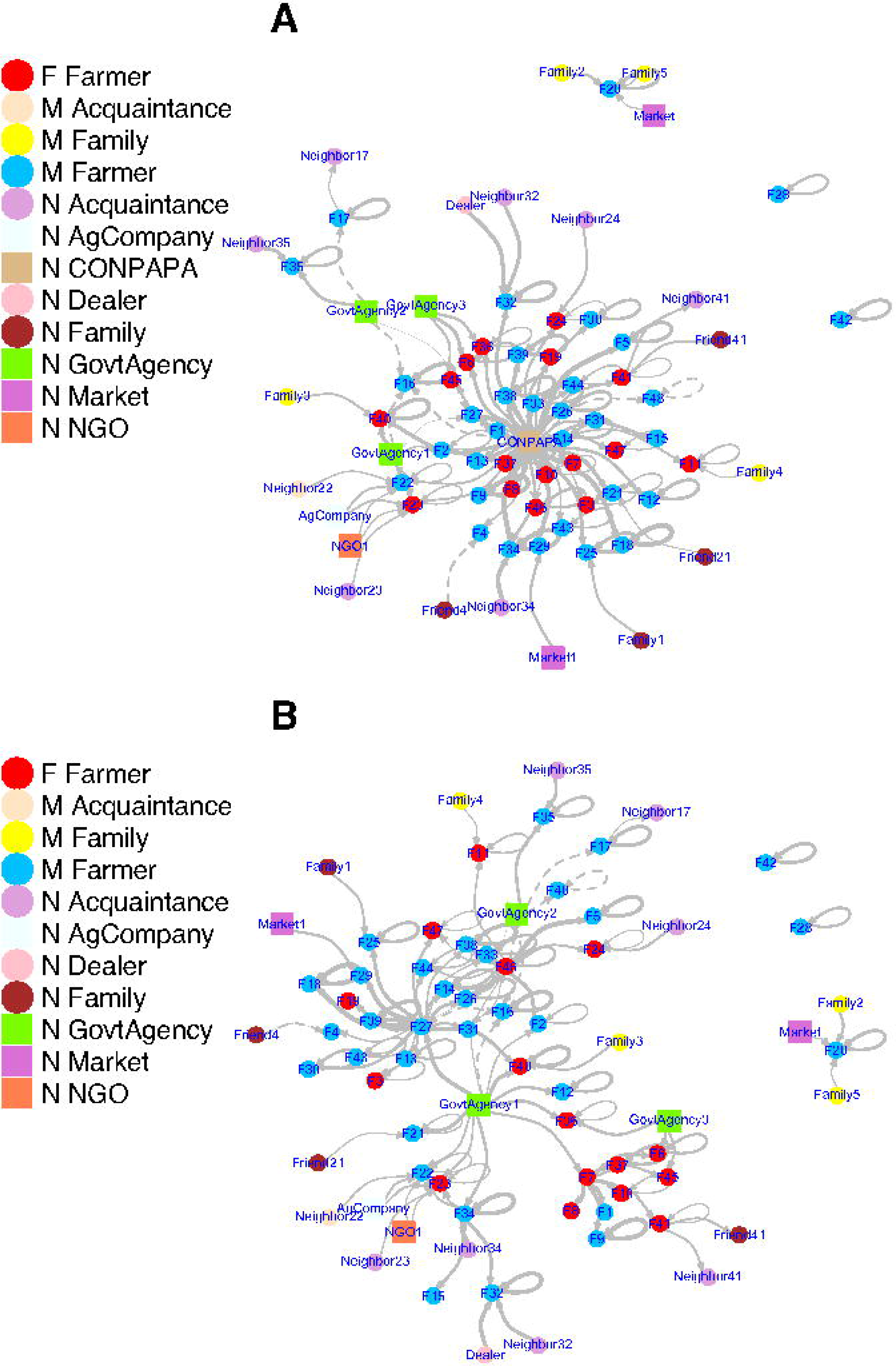
A scenario analysis evaluating potential compensation in the system if the CONPAPA management team no longer played its central role. The figure compares the current scenario, where it provides the majority of seed (**A**), versus a hypothetical scenario where farmers get their seed from the nearest seed multiplier (**B**), and the CONPAPA management team no longer plays a role. **A**, Seed transactions weighted by the volume based on reports from the last three plantings, including CONPAPA. **B**, Seed transactions (links) weighted by volume in a scenario where seed normally going from the CONPAPA management team to multipliers was replaced with seed from the government agency (INIAP). Seed that went from the CONPAPA management team to farmers is now provided by the nearest multiplier. Active multipliers are farmers 7, 27, 34 and 46. For A and B respectively, diameter=4 and 3, density=0.52 and 0.52, and mean of all the shortest paths= 2.1 and 1.4.

## DISCUSSION

In this analysis, we demonstrate an approach for identifying priorities for monitoring plant health in seed systems. In this relatively small and centralized seed system, disease monitoring at CONPAPA processing facilities is obviously a high priority for detection of incipient disease, because it receives high quantities of ware potato (it has high in-degree), and is the source of most of the improved seed (it has high out-degree). Monitoring at the market in Ambato (Market1 in the Figs. 2, 4 and 5) could also be relatively effective. Secondarily, the analysis identifies other nodes in the network that can play a role in sampling and mitigation (Fig. 4), offering a method to rank and prioritize among these nodes for sampling in the field and postharvest. Mitigation measures during a disease outbreak – such as dissemination of new resistant cultivars, training, or treatment of fields – might also prioritize these nodes in the network. Network models provide a window into the epidemiology of plant diseases and strategies for efficient sampling for plant epidemic surveillance and other mitigation efforts (Chadès et al. 2011; Harwood et al. 2009; Hernandez Nopsa et al. 2015; Sanatkar et al. 2015; Sutrave et al. 2012).

Information about the dispersal of particular cultivars through the seed network can provide insights into the likelihood of disease transmission, if cultivars have resistance to a particular disease or if seed of a new cultivar is inadvertently a source of an introduced pathogen. In this simple system, the second most common variety Fripapa was only transacted by 16 of the farmers, so inadvertent spread of a disease in this variety would be far less consequential than in Superchola, which is cultivated by 31 farmers. Good information is available about cultivar susceptibility to *Phytophthora infestans* (e.g., Forbes 2012; Kromann et al. 2009), but studies of viral infection rates for cultivars used in Ecuador rarely consider more than a few varieties. Seed born viral incidence, especially PYVV, PVS, and PVX were reported in one study for some of the cultivars used by CONPAPA farmers, (from lowest to highest incidence: Fripapa, Gabriela, Yana, Unica, Dolores and Chaucha), but per plant yield effects were negligible (Navarrete et al. 2017). High levels of PVY infection have been reported occasionally in Ecuador for Superchola and Fripapa, but viral incidence seems to depend on complex interactions between ecological conditions, on-farm management practices, vector biology, seed sources and cultivar (Navarrete et al. 2017). Yana was reported as extremely resistant to PLRV and PVY, while Unica was resistant to PVY but susceptible to PLRV (CIP 2009).

In Ecuador, seed degeneration, mostly attributable to viruses, can have important effects on yield (7-17% loss, or even *gains* in the case of PYVV), but virus incidence is often low at high altitude, even if levels vary widely from site to site (Peter Kromann unpublished; Devaux et al. 2010; Navarrete et al. 2017; Panchi et al. 2012). It appears that the problem is still under-appreciated and rarely recognized by farmers. For example, only one farmer reported viruses as a concern in this study. Yield losses of ±30% from seed degeneration are common elsewhere in the world (Thomas-Sharma et al. 2017). In most Ecuadorian farms at high altitude it is likely that the vector based transmission rates are lower compared to other seed tuber producing areas worldwide.

We modelled the risk of disease entry into a seed network as a function of farmer information quality and quantity with respect to IPM. This is one approach to integrate the network for the spread of information about management with the biophysical network. A large share of farmers report that they draw on advice from agrochemical stores. Importantly, and perhaps with good reason, farmers do not report trusting them highly as a source of information compared to technical staff working for CONPAPA. Clearly training these store owners about disease and pest management has the potential to be an effective measure to improve management outcomes for farmers inside and outside of the consortium. However, it is unclear if training store owners would result in improved advice and the sale of appropriate pesticides, or if potential economic conflicts of interest would influence the quality of their advice. There may be a financial incentive for small agrochemical store owners to simply recommend the application of the products that they have available for sale.

Women made up a third of the farmers and reported selling smaller volumes of potato product on average. Clearly, they are making less money from potato farming than their male counterparts. There were limited differences in gender access in terms of the number of information sources, or the trust they placed in their information sources. A next step for understanding the role of gender in Andean seed systems would be to determine if this is typical, or if less formal seed system networks in the region reveal larger gender effects. It will also be useful to better understand the potential sources of bias in reporting of trust and other factors; e.g., how does gender influence whether people will tend to report higher or lower levels of trust? An ERGM was used to evaluate the effect of node type on the probability of the existence of reported links. There is great potential for more extensive application of ERGMs in plant disease epidemiology, to test for treatment effects and to estimate and define network structures for applications such as scenario analysis. (For further exploration of this data set using ERGMs, see links at http://www.garrettlab.com/epid-seed/.)

Modeling disease spread in seed and potato transaction networks can indicate the structural effects of seed degeneration. When the value of the first couple of information sources is weighted heavily with diminishing returns for additional information sources, a wider range of risk types is observed among nodes. However, if we assumed that one source would provide good information and new sources incrementally more, the risk associated with most nodes was homogeneous. This highlights the importance of understanding better how farmers use information, and the value of information that farmers receive from different sources. Either scenario is possible. For some diseases, information about a simple IPM intervention could have a large impact, while in other cases, IPM interventions may be complicated to understand or implement, or information about interventions may be poor. Further analysis of this type of system could also integrate the effects of stakeholder knowledge for other components of the network, such as the risk of allowing establishment of disease from seed from other members or the network, or the risk of spreading infected seed to other members of the network.

In the case of viruses, most are transmitted to daughter tubers and will be hitchhikers for each transaction of seed or potato. It is clear that some spread can always occur via the seed system. Network dynamics change from year to year, so scenarios should consider temporal dynamics (e.g., the different effects of wet and dry years; Violon et al. 2016). A more nuanced approach would also take into account different suites of viruses, and the way their transmission rates from infected mother plants to daughter tubers vary depending on varietal and environmental conditions (Bertschinger et al. 2017). Thus node (farmer) vulnerability to infection could also be modeled in terms of specific diseases and scenarios, and could account for varietal differences in resistance. In this system, most transactions were with CONPAPA, governmental institutions or markets in nearby towns, mostly in Ambato which was 10 km from all the member farmers. Only 7% of transactions were with nearby neighbors (1% with family), and always of low volume. In many systems, the probability of transactions or epidemic spread between two cities or organizations follows a gravity model, i.e., it is a function of the distance and the product of the size of the two entities (Jongejans et al. 2015).

A key point to consider for potato seed systems is virus transmission mechanisms. As a case in point, *Potato virus X* (PVX) and *Andean potato mottle comovirus* (APMoV) are transmitted by contact while others such as *Potato virus Y* (PVY) and *Potato leafroll virus* (PLRV), *Potato yellow vein virus* (PYVV) are vectored by aphids (Fankhauser 2000). Networks could include both spread through seed transactions, and spread based on the spatial proximity of farm pairs (as a proxy for the probability of vector movement between a pair). In this study, farms were widely dispersed with both potato and other crops being grown in the intervening areas. Inoculum sources could come from non-CONPAPA potato farmers, or non-potato host species. To realistically model seed infection by vectors would require detailed disease specific data sets that support accurate estimation of dispersal kernels, including the effects of infected volunteers and tuber waste from potato harvesting.

Implementation of fully formal seed systems in many low-income countries is beyond the available resources of the agencies and farmers involved. Reaching the quality levels indicated in statutes may not be feasible. This means that most potato farmers in low-income countries operate wholly within informal seed systems. The CONPAPA seed system has been described as a mixed formal and informal system (Kromann et al. 2017). CONPAPA defines seed quality explicitly in three levels with real quality control measures in place. This means farmers can buy improved seed of known quality with achievable quality levels for the stakeholders involved. The adoption of this alternative seed quality assessment scheme has been incorporated into formal Ecuadorian seed regulation (Kromann et al. 2017), thus formalizing the standards CONPAPA developed. This has been described as “providing flexibility” (FAO 2006) and is recommended as a means of achieving greater confidence by stakeholders and greater adoption of improved seed. Therefore, the CONPAPA seed system could be characterized as predominantly formal with the quality declared seed sources accounting for 47% of the seed in this study. In practice, the mean time between seed replenishment was reported to be approximately 3-4 seasons, though we also found that improved seed is often planted together with reused seed in any given year. This is a much higher rate of improved seed use than the 2-3% formal seed sources reported for Ecuador and Bolivia (Almekinders et al. 2007; Devaux et al. 2010). There are multiple farmer cooperatives that follow the CONPAPA model in central Ecuador and all are adding value. A local-leader at Tungurahua has helped to achieve particularly high levels of cohesion, and provide tangible benefits to the member farmers there. CONPAPA’s cooperative model, combined with the seed quality assessment system could help to overcome issues of access and household economic insecurity that determined participation in formal seed systems elsewhere (Okello et al. 2016). This could have important consequences since potato is becoming increasingly important as a staple crop in areas where informal seed systems prevail (Devaux et al. 2014).

We evaluated the CONPAPA structure as a first step to support improved sampling, IPM, risk assessment for pathogen and pest movement, and farmer decision-making. Identification of key control points that influence the success of seed systems (e.g., farmers, farms, information sources) supports enhancement of the system (e.g., maximizing the distribution of new seed varieties using fewer distribution channels, managing disease outbreaks, and targeting improvement of communication and infrastructure). Resources can be invested in particular nodes to improve practices to control pest and disease outbreaks, leading to improvements in the seed system. We present results for the CONPAPA system as part of an ongoing project to develop general recommendations for improving seed system structure. While we illustrate here how a seed system could *potentially* be resilient to removal of a key node (Fig. 5), the temporal and structural dynamics of seed systems such as CONPAPA need to be better understood to anticipate how they will react to important stressors, and to develop strategies for reducing disease risk while increasing availability of improved varieties.

Specific network configurations are known to influence the probability of disease transmission and persistence (Moslonka-Lefebvre et al. 2011) with the potential for important effects in plant trade networks (Pautasso et al. 2010). The network studied here has small-world properties in that links to and from the CONPAPA management team provide shortcuts across the network. Consistent with scale-free networks, in which nodes are preferentially connected to already highly connected nodes, the CONPAPA management team and the Ambato market act as important hubs. Small-world and scale-free network structures may provide efficient spread of varieties, but also may have high epidemic risk (Moslonka-Lefebvre et al. 2011). Long distance links with the CONPAPA management team indicate the high risk of diseases entering the system from multipliers that provide seed to CONPAPA. Disease management should begin with a focus on them and on the CONPAPA facilities. It is unclear to what extent diseased ware potato could contaminate seed at these facilities. Presumably contamination could occur for some bacterial or fungal pathogens, while being less important for viral pathogens which are the main cause of seed degeneration.

Seed systems share many traits with other managed ecological systems in which there are larger-scale human institutions driving some system components (e.g., federal policy makers and federal research laboratories) and individual land managers who make choices about smaller units in the landscape (e.g., farmers or conservation managers). The approach to scenario analysis presented here can be applied to broader systems that include seed production. A fuller understanding of epidemiological risk may be gained by integrating the risk components evaluated here, due to the network structure of seed transactions and communication, with other risk components such as weather and vector movement. Local seed systems such as CONPAPA are linked internationally via plant breeding networks, through which resistance genes may be distributed (Garrett et al. 2017), with the associated need to manage connectivity for potential movement of pathogens along with germplasm. Local and regional network configurations can also determine the persistence and spread of different cultivars in the landscape, impacting farmer access to genetic resources that may be needed to respond to emerging diseases (Pautasso et al. 2013). Linking epidemiological risk assessment in local seed systems with global seed and germplasm exchange offers an opportunity to expand conceptual frameworks in epidemiology, and to integrate epidemiological concepts with other global risk factors that influence crop yield gaps. Understanding at a systems level can also inform institutional interventions via policy, training, funding or direct management.

## ACKNOWLEDGEMENTS

This work received institutional review board approval through the University of Florida IRB-02 #IRB201700024. We confirm farmer participation was voluntary and personal and demographic information was protected. We appreciate input from Phytopathology reviewers for improving the manuscript. This research was undertaken as part of, and funded by, the CGIAR Research Program on Roots, Tubers and Bananas (RTB) and supported by CGIAR Fund Donors http://www.cgiar.org/about-us/governing-2010-june-2016/cgiar-fund/fund-donors-2/, USDA APHIS grant 11-8453-1483-CA, US NSF Grant EF-0525712 as part of the joint NSF-NIH Ecology of Infectious Disease program, US NSF Grant DEB-0516046, and the University of Florida. CEB wrote R scripts, performed analyses, produced the figures, and wrote the paper; JHN designed the survey, collected data, wrote R scripts, performed preliminary analyses, and wrote the paper; KFA contributed to R scripts and analyses; PK, GAF, and JAP designed the study, and helped collect data; STS contributed to survey design and wrote the paper; PU contributed to survey design and provided an economic analysis; and KAG designed the study, wrote R scripts, mentored authors, and wrote the paper. We appreciate the help of J. L. Brisbane and S. L. Lei in data management, and the help of I. Navarrete in testing preliminary survey drafts. The authors confirm they have no conflicts of interest.

